# Microtubule-templated actin assembly by septin9 drives apical expansion of epithelial cells

**DOI:** 10.64898/2026.05.17.725777

**Authors:** Angelo Arrigo, Victoria Hua, Elias T. Spiliotis, Saurabh Kulkarni

## Abstract

Cells dynamically regulate the size of their apical domain during epithelial morphogenesis. Multiciliated cells (MCCs) represent an extreme case, undergoing massive apical expansion to accommodate hundreds of basal bodies that generate an array of motile cilia. While actin-generated forces drive this expansion, the molecular mechanisms that govern actin assembly and organization remain poorly understood. We show that in *Xenopus*, depletion of Septin9 results in failure of filamentous actin assembly, apical domain expansion, and ciliogenesis. Notably, human Septin9 (SEPT9) localizes to cortical microtubules rather than actin filaments. Live-cell imaging and pharmacological studies reveal that SEPT9 and microtubules form a mutually stabilizing scaffold that directs actin assembly. Using isoform-specific depletion and rescue experiments that genetically separate the microtubule from its actin-bound functions of SEPT9, we demonstrate that SEPT9 association with microtubules is required for actin assembly. These results provide the first in vivo demonstration that microtubule-templated actin assembly via Septin9 drives apical surface expansion during epithelial morphogenesis.

## INTRODUCTION

Cells dynamically regulate the size of their apical domain throughout epithelial morphogenesis. Constriction of the apical surface facilitates processes such as neural tube closure(1, 2), lens placode invagination(3), and gastrulation(4, 5), while expansion contributes to the formation of complex apical architectures observed in vertebrate multiciliated cells (MCCs)(6-8). Although apical constriction has been extensively characterized at the molecular level, ranging from Shroom-mediated actomyosin recruitment to RhoA/ROCK-driven contractility(2, 9-11), the mechanisms underlying apical expansion remain much less well understood(6, 7, 12).

MCCs represent the most extreme case of apical expansion observed in vertebrate development(6, 7). During differentiation, MCCs insert into the outer epithelium and massively expand their apical surface. They establish a dense apicomedial actin meshwork that provides the force necessary for expansion and facilitates the docking of hundreds of basal bodies, each serving as a template for a motile cilium(6-8). Concurrently, a cortical microtubule lattice forms, connecting neighboring basal bodies to establish ciliary polarity(13, 14). Both actin and microtubule networks are assembled de novo at the apical surface during this process, and failure of MCC maturation causes motile ciliopathies, including primary ciliary dyskinesia, infertility, and hydrocephalus(15, 16). Yet how the assembly of these two networks is coordinated, and whether one instructs the other, remains a central unresolved question.

Coordination between actin and microtubule networks is fundamental to cellular function(17, 18). Cells polarize, migrate, divide, and build specialized architectures by coupling these two cytoskeletal systems. Multiple classes of proteins that crosslink actin filaments and microtubules have been identified, but their activities have been characterized largely through in vitro reconstitution and cell culture experiments(19-23). The few in vivo studies of these crosslinkers address cell migration and wound healing in the mouse epidermis(24),and axon extension and tube formation in Drosophila(5, 25).

Septins constitute a family of GTP-binding proteins that polymerize into nonpolar filaments and ring structures and are increasingly recognized as regulators of actin and microtubule organization(26). Specifically, the septin complex has been shown to crosslink networks and template actin polymerization on the MT lattice(27). Septins have been localized to the basal bodies and axonemes of primary cilia(28-31). SEPTIN2 (SEPT2) forms a diffusion barrier at the base of the cilium that establishes and maintains ciliary membrane protein distribution(30). SEPT9 binds and activates the RhoGEF ARHGEF18 to recruit the exocyst complex during transition zone assembly(31). Septin complexes along the axoneme regulate ciliary length, in part by stabilizing the axonemal microtubules(29, 32). In contrast, septin function in MCCs has been examined by only two studies, with one study largely focused on the role of SEPT7 in planar cell polarity signaling in MCCs(33) , and a second exploring the localization of septins in the cilia of airway epithelial cells(34).

Here, we sought to investigate how septins coordinate the actin and microtubule cytoskeletons during MCC morphogenesis. We focused on Septin 9 (*SEPTIN9* in humans, *septin9* in *Xenopus*) because it directly interacts with both actin filaments and microtubules via the basic domain of its N-terminal extension, which is unique in the septin family(26, 27, 35). In vitro studies have shown that SEPT9 binds to and cross-links actin stress fibers, stabilizes focal adhesions, and inhibits cofilin-mediated actin depolymerization(26, 35-37). Its positively charged residues facilitate electrostatic interactions with acidic beta-tubulin, leading to microtubule bundling(26, 27, 35, 38-40). Notably, septin complexes can directly crosslink microtubules and actin filaments into hybrid bundles and template actin polymerization on microtubule lattices(41), and SEPT9 specifically binds and bundles microtubules through its N-terminus(26, 35). This suggests that SEPT9 might serve as a molecular link between these two cytoskeletal networks that MCCs need to coordinate. However, it remains unknown whether SEPT9 coordinates actin and microtubule remodeling during development in vivo.

Using *Xenopus tropicalis*, we demonstrate that Septin9 is critical for the expansion of the MCC apical surface by regulating F-actin assembly. However, Septin9 localizes to cortical microtubules rather than the F-actin network it is required to build. Pharmacological disruption of the cytoskeleton and live-imaging studies show that Septin9 and microtubules create a mutually stabilizing scaffold that guides actin assembly. Structure-function analyses identify a domain hierarchy that determines cytoskeletal network selection, and genetic experiments indicate that Septin9 and microtubule association is necessary for complete apical expansion, whereas its actin-binding capability alone is not sufficient. These results provide in vivo evidence that microtubule-to-actin remodeling by Septin9 drives apical surface morphogenesis.

## RESULTS

### Septin9 is essential for apical expansion and F-actin organization in multiciliated cells

To investigate the role of Septin9 in MCC development, we depleted the long isoform of *septin9* (XM_031894339.1, *septin9-x1*) in *Xenopus tropicalis* embryos using a translation-blocking morpholino and evaluated MCC function at mid-tailbud stages (stage 28), when the ectoderm is covered with mature MCCs. Coordinated ciliary beating generates anteroposterior fluid flow across the ectoderm, serving as a functional indicator of MCC maturation (Figure 1a). Microbead flow assays showed that control embryos cleared beads at approximately 202 µm/sec, while *septin9-x1* morphants exhibited significantly impaired clearance (∼27 µm/sec) (Figure 1a–a’, c).

**Figure 1:**
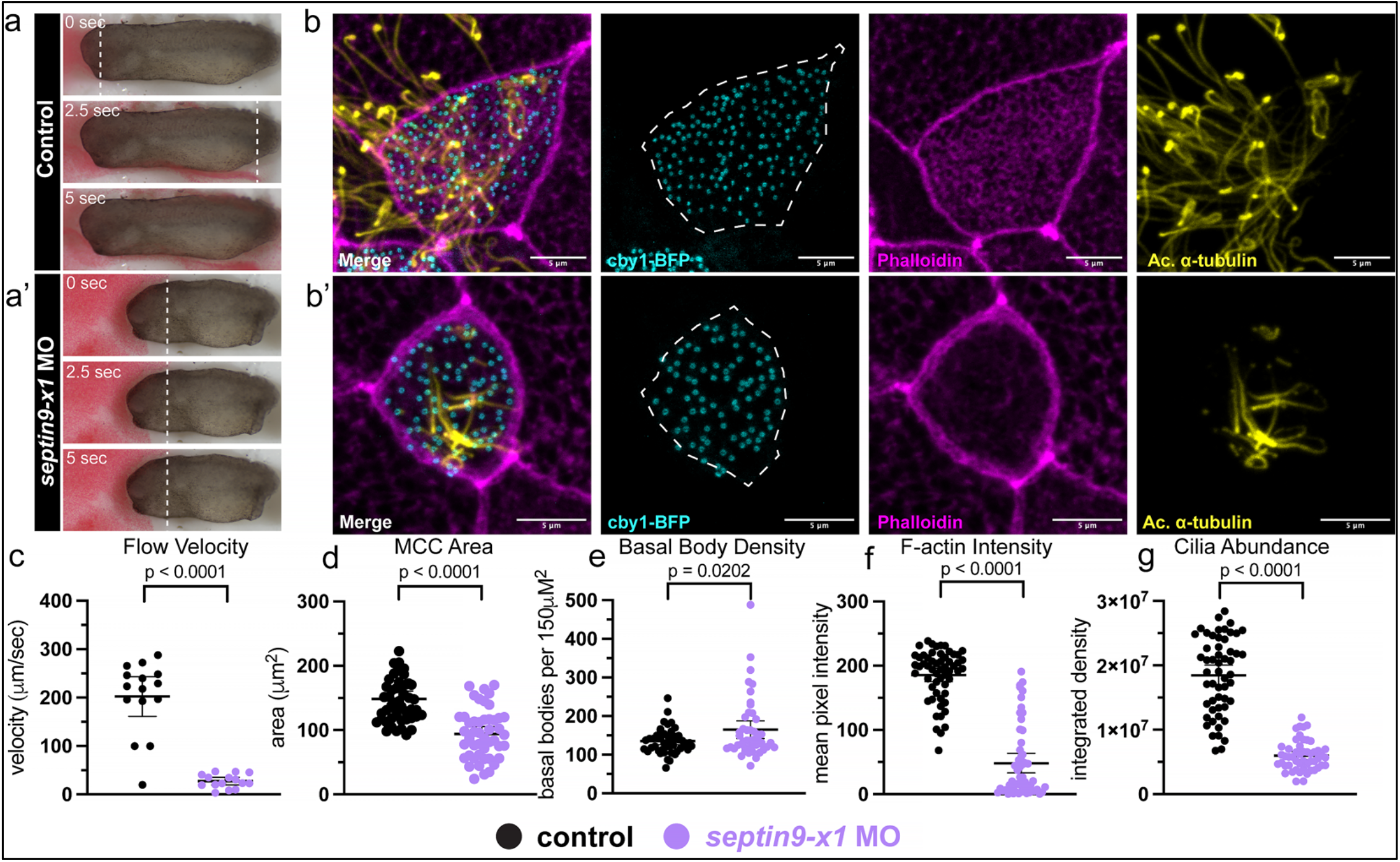
Septin9 is essential for apical expansion, F-actin organization, and ciliogenesis in multiciliated cells. **(a-a’)** Microbead flow assay shown at timepoints 0, 2.5 seconds, and 5 seconds of both an injected control *Xenopus tropicalis* embryo and a *septin9-x1* morphant embryo at stage 28. The dashed white line represents the front of the leading beads in each still. **(b-b’)** Confocal max intensity projections from acetylated *α*-tubulin (cilia, yellow) and phalloidin (F-actin, magenta) staining cby1-BFP (basal bodies, cyan) injected control and *septin9-x1* morphant embryos at stage 28. **(c)** Quantification from panels a and a’ showing bead flow velocity. **(d-g)** Quantification of MCC area, basal body density, MCC F-actin intensity, and cilia abundance as determined using Fiji software. For flow velocity: n=15 embryos per condition; for MCC area, basal body density, MCC F-actin intensity, and cilia abundance: n=12 embryos and 47-54 cells per condition. Scale bar is 5µm for each IF image.

Immunofluorescence analysis revealed the cellular basis of this functional impairment. In control MCCs, a dense apicomedial F-actin meshwork covered the apical surface, forming a continuous lattice that provides structural support for apical expansion, basal body docking, and spacing to facilitate motile cilia formation (Figure 1b). In *septin9-x1* morphants, this F-actin network was absent (Figure 1b, b’, f). Without this scaffold, the apical domain failed to expand; morphant MCCs were circular rather than polygonal and showed a marked reduction in area compared with controls (Figure 1b, b’, d). The lack of actin and the failure to expand directly affected basal body number and distribution: while basal bodies were uniformly spaced (nearest neighbor distance) in both controls and morphants, they appeared clustered near the cell center in morphant MCCs (Figure 1b, b’, e, S1). Correspondingly, cilia abundance was significantly affected in morphants (Figure 1b’, g). In summary, Septin9 depletion disrupts the entire apical maturation process: the F-actin network fails to assemble, the apical domain does not expand, basal body organization is compromised, and ciliogenesis is impaired.

### Septin9 is essential for the de novo assembly of the apical actin network during apical expansion

The phenotype suggested that septin 9 (SEPT9) either facilitates the assembly of an F-actin network or is critical for maintaining its assembly. To distinguish between these two possibilities, MCCs were examined at four developmental stages representing different stages of MCC maturation.

First, we examined the progression of MCC maturation, comparing control and morphant embryos across four developmental stages. In controls, MCCs emerged at the apical surface by stage 18 with a small apical domain and no mature F-actin meshwork (Figure 2a). By stage 22, an organized F-actin network had begun to assemble alongside apical expansion, and through stages 25 and 28, this network developed into a dense, intricate meshwork while the apical domain continued to grow (Figure 2a, b–c). This developmental progression was markedly altered in morphants. At stages 18 and 22, fewer morphant MCCs had reached the apical surface, indicating delayed emergence (Figure 2a, d). By stage 25, morphant MCC numbers had caught up with controls (Figure 2d), demonstrating that *septin9-x1* is not required for MCC specification or radial intercalation itself, but only for the timely completion of apical expansion. However, even after morphant MCCs reached the surface, the apical F-actin network was never assembled, as evidenced by the lack of F-actin enrichment at the apical surface (white square insets Figure 2a, a’). Moreover, the apical domain never expanded - the apical area remained minimal through stages 25 and 28 (Figure 2a, b–c). This temporal separation reveals two distinct requirements for SEPT9 in MCC morphogenesis: first, for the timely radial intercalation of nascent MCCs into the outer epithelium, and second, and more dramatically, for the post-emergence assembly of the apical F-actin network that drives apical expansion.

**Figure 2:**
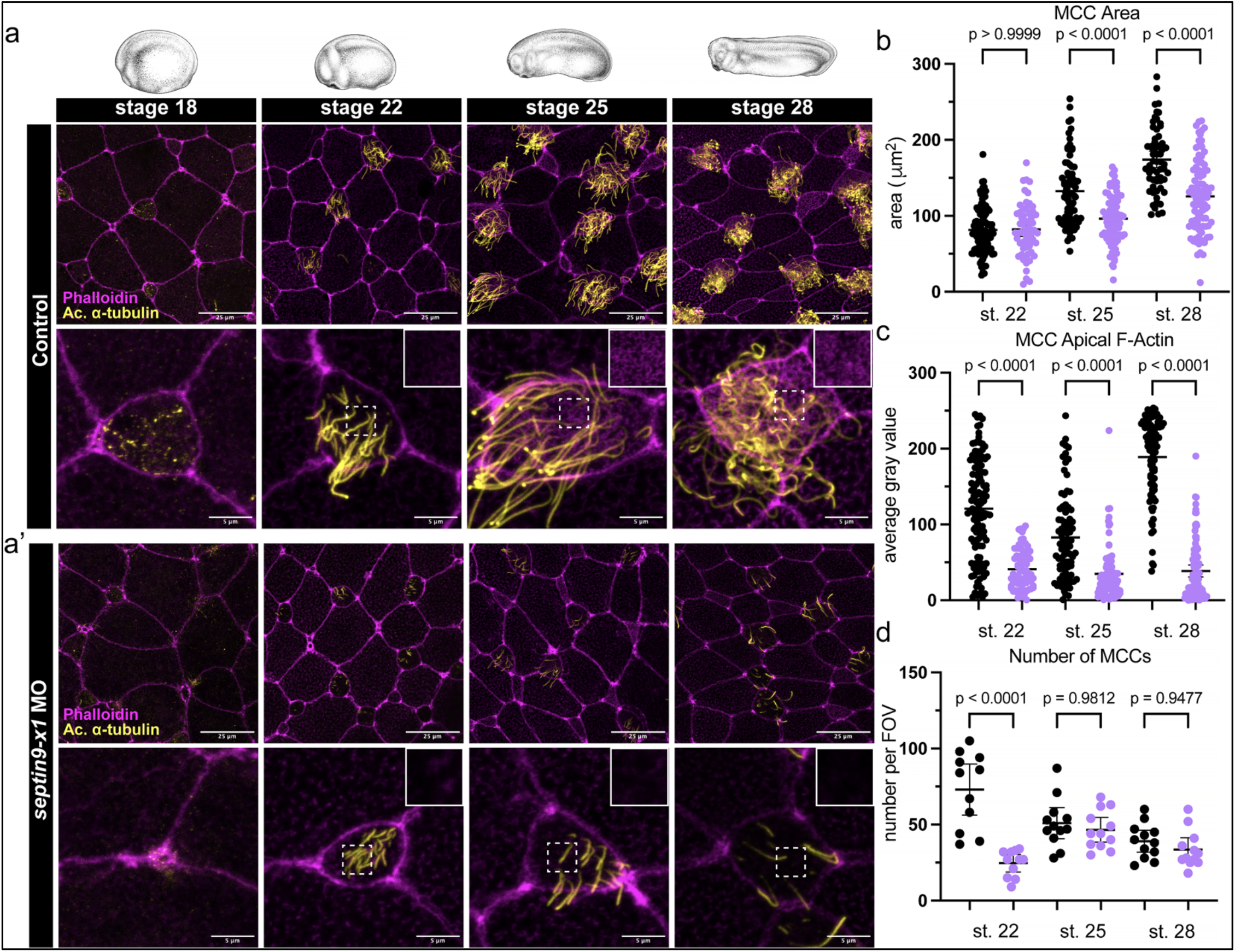
Septin9 is essential for the de novo assembly of the apical actin network during apical expansion in multiciliated cells. **(a-a’)** Drawings of whole embryos at stage 18, stage 22, stage 25, and stage 28, and representative confocal max intensity projections from controls and *septin9-x1* morphants at each stated timepoint, stained with both acetylated *α*-tubulin (cilia, yellow) and phalloidin (F-actin, magenta). Both low (2x) and high (6x) zoom images are shown. Insets marked by white squares show F-actin enrichment at the apical surface of respective MCCs. **(b)**Quantification of MCC area at stages 22, 25, and 28. **(c)**Quantification of MCC F-actin intensity at stages 22, 25, and 28. **(d)**Quantification of the number of MCCs within the field of view at stages 22, 25, and 28. Stage 18 is excluded because MCCs had not yet fully emerged at that stage. For MCC area and MCC F-actin intensity: n= 12 embryos and 59-123 cells per condition. For the number of MCCs: n=11-12 embryos per condition. Scale bar is 25µm for each low-zoom image and 5µm for each high-zoom image.

### SEPT9 localizes to cortical microtubules, not to the F-actin network via its microtubule binding domain

Having established that Septin9 is crucial for actin assembly and apical expansion, we asked where SEPT9 localizes in MCCs. Given its requirement for the actin network, we hypothesized that SEPT9 would colocalize with F-actin and directly cross-link actin filaments to form a meshwork. To test this, we expressed human SEPT9_i1 (NP_001106963.1, orthologous to *X. tropicalis septin9-x1*) fused to BFP (SEPTIN9-i1-BFP) in *Xenopus* MCCs and co-imaged it with the F-actin marker utrophin (RFP-UTRN) (see Fig S2 for all localization images). Surprisingly, SEPT9_i1 did not colocalize with apical F-actin. Instead, it occupied the gaps between actin filaments, displaying a pattern similar to the cortical microtubule network (Figure 3a). We therefore examined its relationship with cortical microtubules, which were labeled with MAP7-GFP, and found that SEPT9_i1 colocalizes with the cortical microtubule lattice within the apical plane (Figure 3b). This result presented a paradox: depletion of Septin9 leads to loss of the apical actin network, but SEPT9_i1 colocalizes with cortical microtubules.

**Figure 3:**
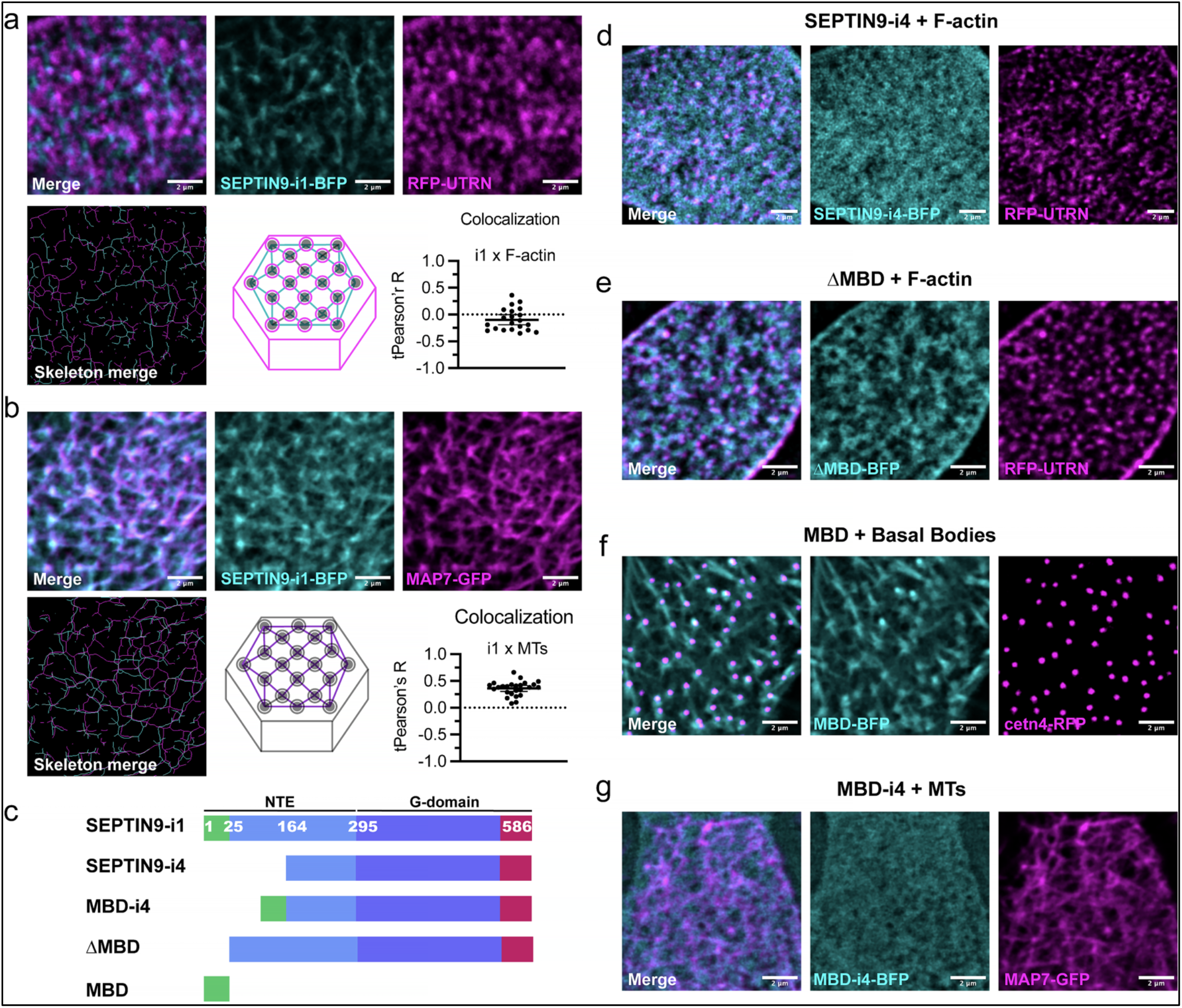
Septin9 localizes to cortical microtubules, not to the F-actin network via its microtubule binding domain (MBD, 1-25 aa). **(a)**Live image of SEPT9_i1-BFP (cyan) localization and RFP-UTRN (F-actin, magenta) localization in an injected *X. tropicalis* embryo. This same image was skeletonized using a Fiji script. Colocalization was determined between SEPT9_i1-BFP and RFP-UTRN. **(b)**Live image of SEPT9_i1-BFP localization and MAP7-GFP (microtubules, magenta) localization. This same image was skeletonized using a Fiji script. Colocalization was determined between SEPT9_i1-BFP and MAP7-GFP. **(c)**SEPT9 constructs designed for use in this manuscript. The MBD-i4, ΔMBD, and MBD constructs were derived from Kuzmić et al. 2022(35, 40) . **(d)**Live image of SEPT9_i4-BFP (cyan) localization and RFP-UTRN (F-actin, magenta) localization. **(e)**Live image of ΔMBD (human SEPT9 isoform 1 with the first 25 amino acids deleted, cyan) localization and RFP-UTRN (F-actin, magenta) localization. **(f)**Live image of MBD-BFP (first 25 amino acids of human SEPT9 isoform 1, cyan) localization and cetn4-RFP (basal bodies, magenta) localization. **(g)**Live image of MBD-i4-BFP (first 25 amino acids of human SEPT9 isoform 1 fused to isoform 4, cyan) localization and cetn4-RFP (basal bodies, magenta) localization. Each of these images is a zoomed-in view of one MCC. For colocalization analyses: n=21-26 cells. Each confocal image is a max intensity projection that has been deconvolved (2-5 iterations) using Fiji. Scale bar is 2µm for each image.

To identify the domains responsible for the microtubule localization, we tested a panel of constructs based on structural differences among *SEPT9* isoforms (Figure 3c). Isoform 1 contains a unique N-terminal extension (amino acids 1– 256), including a microtubule-binding domain (MBD; amino acids 1–25), whereas isoform 4 is identical but lacks the basic domain (aa 1-164) of this extension, which has been previously shown to interact directly with microtubules and actin filaments(26, 27, 35, 40). SEPT9_i4 appeared diffuse and lacked specific cytoskeletal association (Figure 3d), indicating the basic domain of the N-terminal extension is essential for localization. Deletion of the N-terminal first 25 amino acids (1-25aa), MBD from isoform 1 (ΔMBD)(35), shifted localization from cortical microtubules to F-actin (Figure 3e), demonstrating that the first 25 aa MBD is necessary for microtubule association and revealing an actin-binding activity normally overridden when the MBD is present. Expressing this MBD alone (amino acids 1–25) resulted in consistent localization to basal bodies and inconsistently to the basal body-associated microtubule-like filaments (Figure 3f), suggesting the MBD provides tubulin-binding capacity but is insufficient for full cortical microtubule association, which may require additional microtubule-binding motifs downstream of the MBD(40). Fusion of the MBD to isoform 4 (MBD-i4) did not produce specific localization (Figure 3g), which might be due to masking or allosteric inhibition of the MBD peptide by the acidic and/or G-domain of the isoform 4 or lack of synergy with the sequences within the basic 1-146 aa domain(35, 40).

### Isoform-specific depletion reveals genetically separable microtubule and F-actin functions

Based on the structure-localization analysis, we hypothesized that isoforms 1 and 4 may exert distinct effects on the MCC cytoskeleton. Isoform 1, which localizes to cortical microtubules, is responsible for organizing both the microtubule and F-actin networks. Conversely, isoform 4, which lacks the MBD and cannot bind microtubules, would primarily regulate F-actin organization. To investigate this, isoform-specific morpholinos were designed, and the cytoskeletal effects of depletion of individual variants were compared.

To deplete each isoform individually, we designed isoform-specific translation-blocking morpholinos that exploit the partitioning of translation start sites by *septin9* alternative splicing. Although *septin9-x1* and *septin9-x5* (human *SEPT9-i4*) share their C-terminal coding sequence encoding the Septin9 core, they initiate translation at different AUG codons: x1 uses an upstream start that produces the unique N-terminal extension (aa 1–164), while x5 uses a downstream start at the beginning of *septin9-x5*. The sequence encoding the x1 N-terminal extension is therefore part of the *x1* mRNA but constitutes the 5′ UTR of *x5* mRNA. Because translation-blocking morpholinos function by sterically blocking the scanning initiation complex and are effective only when bound within a window spanning the 5’ UTR through the first ∼25 nucleotides of the coding sequence(42, 43), each morpholino blocks only the isoform whose AUG it covers. MO-x1 occupies the 5′ UTR/AUG region of x1 and blocks its initiation; in *x5* mRNA, the same sequence lies far upstream of the x5 AUG and is therefore outside the effective window. MO-x5 occupies the AUG region of x5; in *x1* mRNA, this sequence lies deep within the coding region, downstream of the x1 initiation site, where bound morpholinos are displaced by elongating ribosomes

Morphants for *septin9-x1* exhibited disrupted cortical microtubule networks characterized by short, fragmented microtubules and a loss of the typical intersecting lattice pattern in the apical domain (Figure 4a-b). Quantification showed a significant reduction in the size of the largest connected component of the microtubule network (Figure 4e). *Septin9-x1* depletion also resulted in an almost complete loss of apical F-actin, consistent with previous full morpholino experiments (Figure 4a, c). In contrast, *septin9-x5* morphants did not significantly affect the cortical microtubule network and component length (Figures 4a–b, e) but showed significant disruption of apical F-actin meshwork (Figure 4a, c), suggesting that *septin9-x5* specifically contributes to actin organization independently of microtubules.

**Figure 4:**
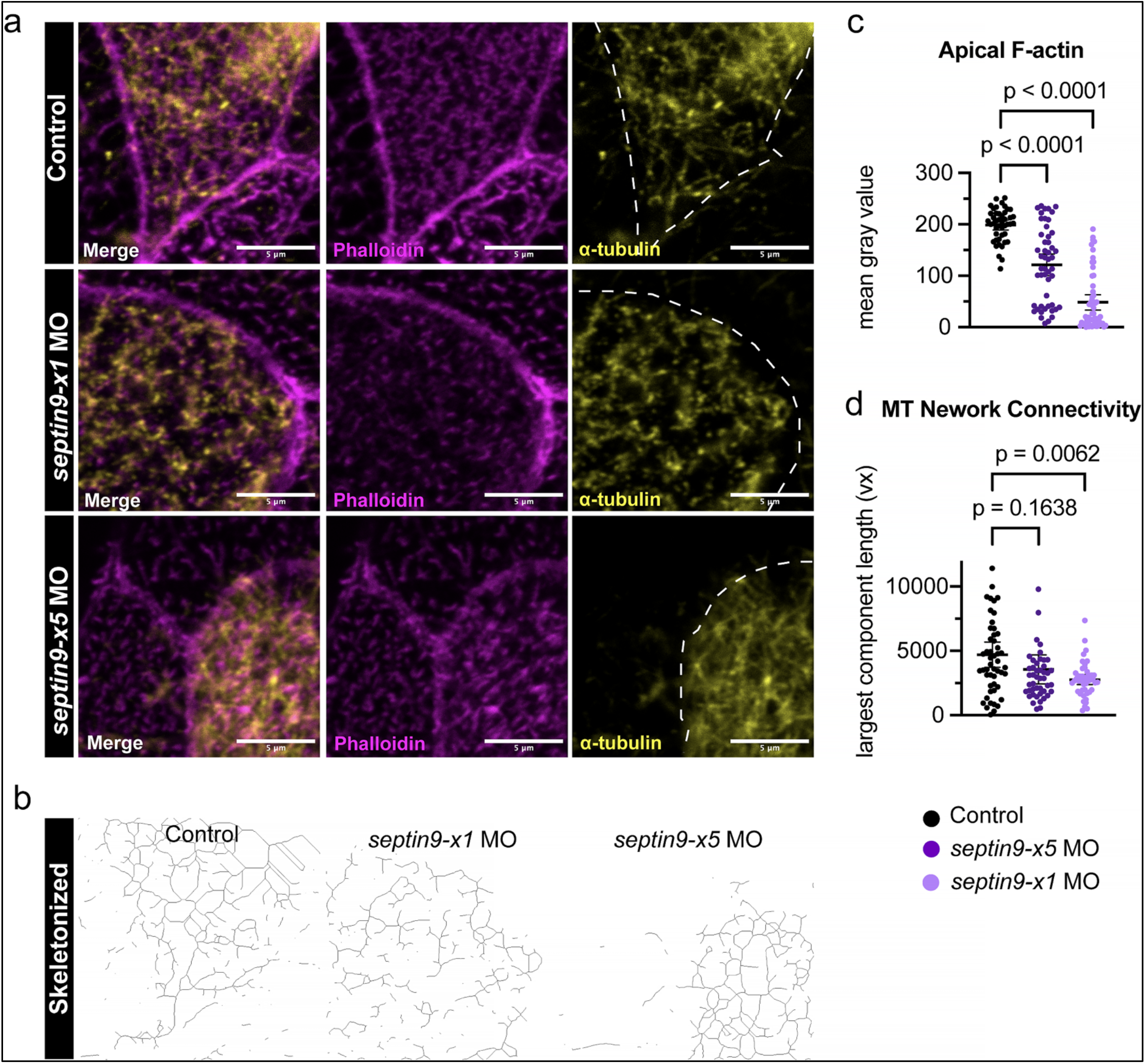
Isoform-specific depletion reveals genetically separable microtubule and F-actin functions. **(a)**Confocal max intensity projections of an injected-control, *septin9-x1* morphant, and *septin9-x5* morphant embryo stained with acetylated *α*-tubulin (cilia, yellow) and phalloidin (F-actin, magenta). The dotted line represents the cell barrier determined by the phalloidin channel. **(b)**Skeletonized images of the microtubule network from panel a. Processing (subtraction and blurring) and skeletonization were performed in Fiji. **(c)**Quantification of the MCC apical F-actin intensity. **(d)**Quantification of the MCC subapical F-actin intensity. **(e)**Quantification of the MCC microtubule network connectivity. All images were analyzed with Fiji software. For MCC apical/subapical F- and network connectivity: n = 12 embryos and 42-54 cells per condition. Scale bar is 5µm for each image.

These findings demonstrate that the microtubule-organizing and actin-promoting functions of Septin9 are genetically separable. Isoform 1, containing the MBD within the N-terminal extension, is essential for cortical microtubule organization, and its loss disrupts both networks. Isoform 4 facilitates actin assembly through a microtubule-independent mechanism, likely via the conserved actin-binding capacity of the SEPT9 domains (GTP-binding and acidic poly-proline) shared by both isoforms.

### Septin9 and cortical microtubules form a mutually stabilizing scaffold that directs actin assembly

The preceding results demonstrate that Septin9 localizes to cortical microtubules and is required for both microtubule organization and apical F-actin assembly, and that these two functions are genetically separable by isoform. However, depleting *septin9-x1* affects both microtubules and F-actin, making the primary dependency unclear. One possibility is that Septin9 uses cortical microtubules as a scaffold, and once localized, stabilizes microtubules and directs actin assembly. Alternatively, F-actin may stabilize Septin9 at the cortex, and Septin9, in turn, organizes the microtubule network. To distinguish between these hypotheses, we combined live imaging with pharmacological disruption of each cytoskeletal system.

To resolve the temporal sequence of network assembly, we co-injected embryos with SEPTIN9-i1-BFP, MAP7-GFP, and RFP-UTRN and imaged emerging MCCs by time-lapse confocal microscopy from stage 18 onward (Figure 5; Figure S3 shows individual channels separately). At t = 15 min, the cortical microtubule network had already begun to organize. SEPT9_i1 appeared as punctate foci, and a few short filaments extending from the puncta were aligned with the cortical microtubule network. F-actin was largely confined to the cell perimeter, but F-actin foci were clearly present on SEPT9_i1 decorated microtubules (MAP7-GFP), and at intersections, therefore, indicative of actin nucleating foci in the absence of a filamentous actin network. By t = 30 min, SEPT9_i1 was more extensively associated with the cortical microtubules as filaments and puncta (Figure 5, insets), and short actin filaments appeared to extend from puncta and co-align with SEPT9_i1 and microtubules. As SEPT9 continued to assemble along the microtubule network, these initial actin nucleation foci progressively expanded into a continuous meshwork by t = 60 min, accompanied by apical domain expansion.

**Figure 5:**
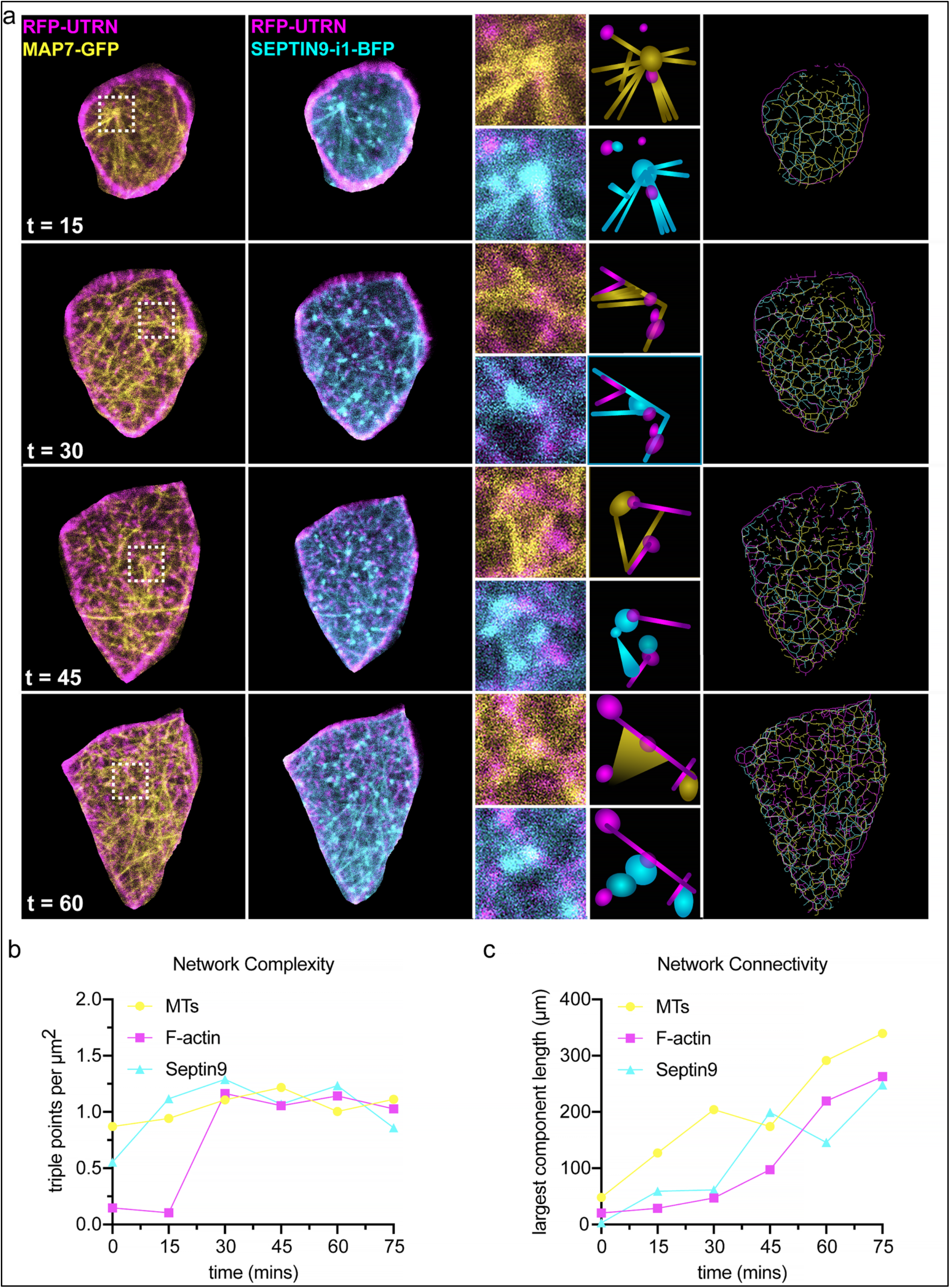
Cortical microtubules template Septin9 organization followed by filamentous actin assembly. **(a)**Time-lapse images (confocal max intensity projections) of a newly intercalated MCC from a stage 18 *X. tropicalis* embryo injected with SEPT9_i1-BFP (cyan), RFP-UTRN (F-actin, magenta), and MAP7-GFP (microtubules, yellow). The same MCC was imaged as it expanded its apical area over 75 minutes. The third column is zoomed images from the left columns, as indicated by the dashed square. A schematic of the interactions between microtubule and F-actin and SEPT9_i1 and F-actin was utilized for column four. Each channel was processed, skeletonized, and merged in the Fiji software to create column five. **(b)**Quantification of the network complexity of the microtubule, F-actin, and SEPT9 networks over time from panel a. **(c)**Quantification of the network connectivity of the microtubule, F-actin, and SEPT9 networks over time from panel a. These graphs are from data extracted from the time-lapse image sequence and therefore represent this MCC’s networks (F-actin, microtubules, and SEPT9) over time.

To quantify these dynamics, we skeletonized each network at successive timepoints and measured both network complexity and connectivity (Figure 5b–c). Microtubule complexity rose first and remained high throughout. SEPT9 complexity rose subsequently, tracking microtubule organization. F-actin complexity rose sharply only after SEPT9 had organized along microtubules, with a clear delay relative to both networks. The connectivity analysis confirmed the same sequence: microtubules reached the connectivity plateau first, SEPT9 second, and F-actin last.

The functional importance of this sequential assembly was confirmed by imaging *septin9-x1* morphant MCCs co-injected with the same fluorescent markers (Figure S4). In contrast to controls, morphant MCCs failed to progress through any stage of this program: the cortical microtubule network never organized into a coherent lattice, and F-actin never accumulated in the apical interior. The apical domain showed no measurable expansion over the 75 minutes imaged. Quantification confirmed that none of the networks gained complexity or connectivity in morphants (Figure S4b–c), in stark contrast to the progressive assembly observed in controls. These data establish that SEPT9 is required not only for the final step of actin assembly, but for the coordinated emergence of the entire cytoskeletal program. Microtubules organize first; SEPT9 follows, and F-actin is built last upon the pre-existing SEPT9/microtubule scaffold. Without SEPT9, the program fails from its earliest stages.

### Pharmacological disruption confirms a directional MT → SEPT9 → F-actin pathway

To test the dependencies between the three networks, we treated embryos expressing SEPTIN9-i1-BFP, MAP7-GFP, and RFP-UTRN with nocodazole (disrupts MTs) or latrunculin A (disrupts F-actin) and assessed network organization by live imaging. In DMSO-treated controls, all three networks displayed their characteristic organization (Figure 6a). Nocodazole treatment eliminated the cortical microtubule lattice and nearly completely abolished the SEPT9 network (Figure 6b), demonstrating that SEPT9 depends on microtubules for its localization. In contrast, latrunculin A disrupted the apical F-actin network but left SEPT9 intact and properly localized on cortical microtubules (Figure 6c).

**Figure 6:**
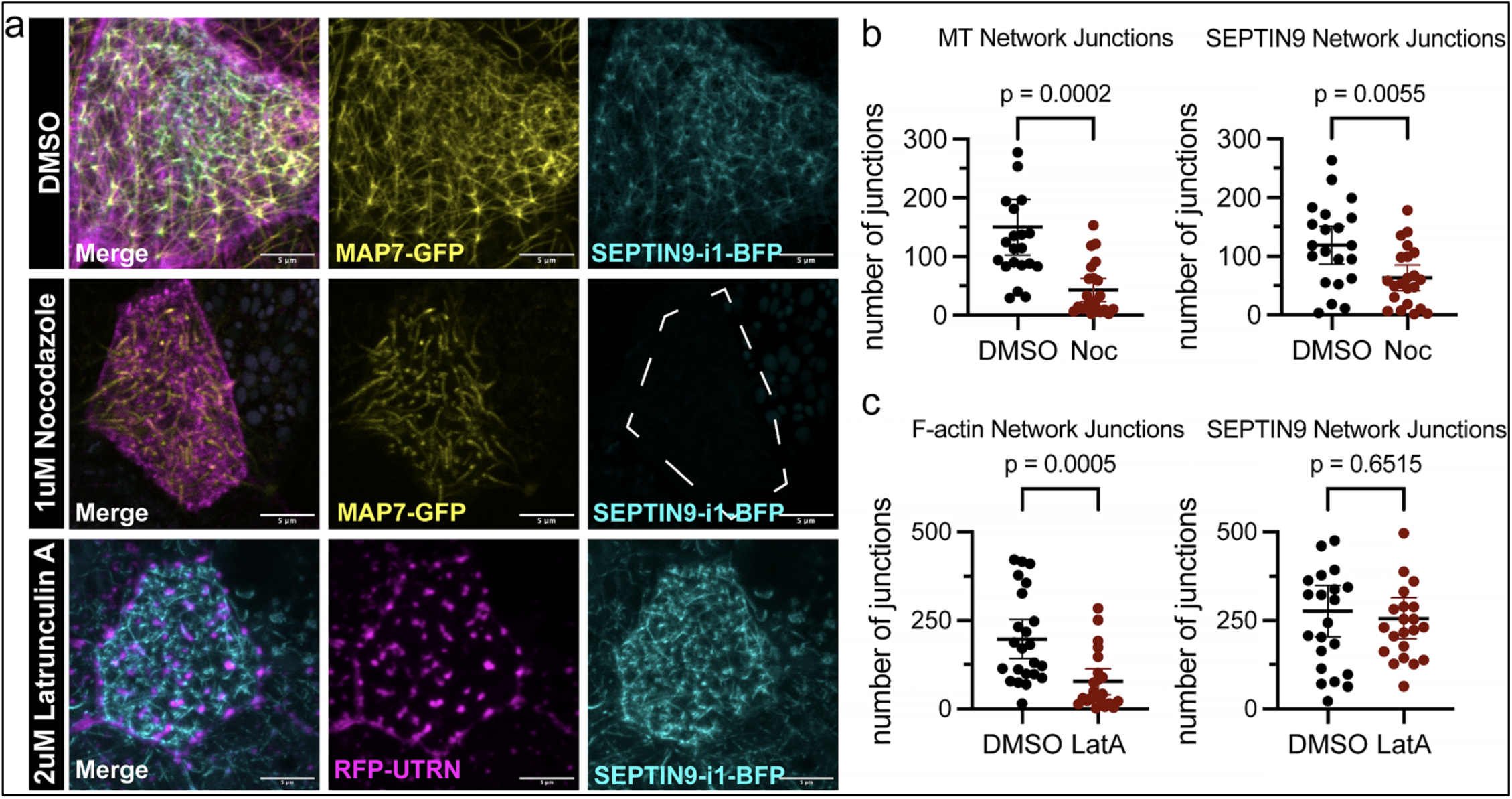
Pharmacological disruption confirms Septin9 depends on cortical microtubule network but not apical F-actin . **(a)**Live images (confocal max intensity projections) of an MCC from an embryo injected with RFP-UTRN (F-actin, magenta), MAP7-GFP (microtubules, yellow), and SEPT9_i1-BFP (cyan) and treated with either DMSO, 1µM nocodazole, or 2µM latrunculin A. Nocodazole treatments were from stage 23-28(13) and imaged at stage 28. Latrunculin A treatment lasted 15 minutes at stage 28(8), and imaging was performed immediately afterward. **(b)**Quantification of the number of junctions from both the microtubule and SEPT9 network in both DMSO and nocodazole-treated embryos. **(c)**Quantification of the number of junctions from both the F-actin and SEPT9 network in both DMSO and latrunculin A-treated embryos. Fiji software was used to process (subtraction and blur), skeletonize, and quantify the number of junctions for each network. For each analysis: n=21-23 cells per condition. Scale bar is 5µm for each image.

No single experiment establishes directionality on its own, but together, the temporal sequence captured by live imaging, the dependency relationships revealed by pharmacology, and the genetic separation of MT and actin functions through isoform-specific depletion converge on a single model. Septin9 and cortical MTs are mutually stabilizing; MTs scaffold Septin9, and Septin9, in turn, organizes the MT network. From this stabilized platform, Septin9 directs the de novo assembly of the apical F-actin network. Actin is the directional output of this scaffold, not a contributor to it.

### Domain-specific rescue confirms that apical expansion requires microtubule-directed Septin9 function

Having established the directionality of the pathway and the domain requirements for SEPT9 localization, we asked whether these domain-specific localizations correspond to distinct functional outputs in apical expansion and actin organization. To test this, we knocked down *septin9-x1* and rescued it using three different mRNAs: full-length SEPTIN9-i1-BFP, SEPTIN9-i4-BFP, and ΔMBD-BFP (Figure 7a, b). Full-length SEPTIN9-i1-BFP rescued MCC apical area, apical F-actin intensity, and, therefore, ciliogenesis (Figure 7b, c). This rescue with the human ortholog demonstrates both that the morpholino specifically targets *septin9-x1* and that SEPT9_i1 is functional in *X. tropicalis* MCCs. SEPT9_i4, which lacks the MBD and cannot localize to cortical microtubules, failed to rescue apical area or ciliogenesis but did rescue apical F-actin intensity (Figure 7b, d). We consider it a partial rescue because the apical F-actin enrichment was insufficient to support apical expansion(6, 12). This result is consistent with isoform 4 retaining partial actin-binding activity while lacking the microtubule-dependent functions required for expansion. ΔMBD, which localizes to F-actin rather than microtubules, phenocopied the isoform 4 rescue with mild restoration of apical F-actin without recovery of apical area or cilia density (Figure 7b, e).

**Figure 7:**
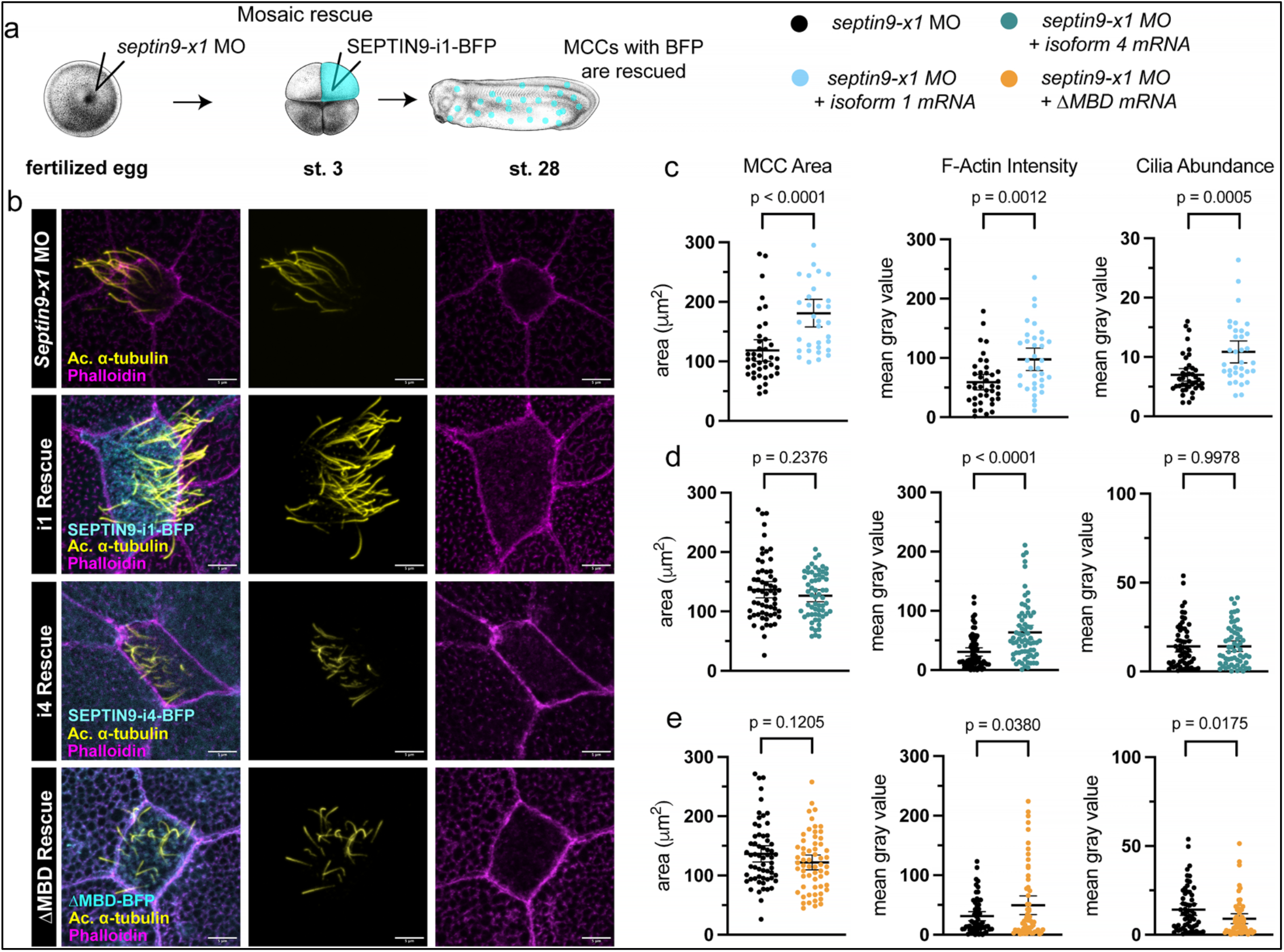
Domain-specific rescue confirms that apical expansion requires microtubule-directed Septin9 function. **(a)**Schematic showing the rescue strategy used for *septin9-x1* morphants with SEPT9_i1-BFP. **(b)** Confocal max intensity projections of a *septin9-x1* morphant and potential rescues: SEPT9_i1-BFP, SEPT9_i4-BFP, and ΔMBD-BFP stained with acetylated *α*-tubulin (cilia, yellow) and phalloidin (F-actin, magenta). The human mRNA injected for each rescue is in cyan. **(c)**Quantification of MCC area, MCC F-actin intensity, and cilia abundance for the SEPT9_i1-BFP rescue. **(d)**Quantification of MCC area, MCC F-actin intensity, and cilia abundance for the SEPT9_i4-BFP rescue. **(e)**Quantification of MCC area, MCC F-actin intensity, and cilia abundance for the ΔMBD-BFP rescue. For each analysis: n=12 embryos and 60-68 cells per condition. Scale bar is 5µm for each image.

The convergence of the isoform 4 and ΔMBD partial-rescue phenotypes is notable: these constructs differ in everything except the absence of the MBD. This confirms that the MBD is the critical determinant of whether rescue is full or partial. Thus, the complete apical maturation program: actin assembly, apical domain expansion, basal body spacing, and ciliogenesis requires SEPT9’s microtubule-directed function through its MBD-containing N-terminal extension.

## DISCUSSION

Our findings identify SEPT9_i1 as the molecular link between the two cytoskeletal networks that build the MCC apical surface: SEPT9 localizes to cortical microtubules and, from this scaffold, directs de novo F-actin assembly (Figure 8). Three convergent lines of evidence establish this directionality. With live imaging, we captured the temporal sequence: microtubules organize first, SEPT9 assembles along them, and F-actin appears 30–45 minutes later upon this pre-existing scaffold. Pharmacology isolates the dependencies, nocodazole eliminates SEPT9, whereas latrunculin A leaves it intact on cortical microtubules. Isoform-specific depletion confirms the genetic hierarchy: loss of the microtubule-binding isoform (isoform 1) abolishes both networks, whereas loss of the non-microtubule-binding isoform (isoform 4) spares microtubules but partially disrupts actin. Taken together, these findings demonstrate the model Microtubule↔SEPT9→Actin, with mutual reinforcement among SEPT9, microtubules, and actin, and with actin as the strictly downstream output. Our data provide the first demonstration that microtubule-to-actin coordination drives apical surface expansion, leading to a morphogenetic outcome in vivo. To our knowledge, this is one of the very few in vivo paradigms of microtubule-based actin assembly, which has emerged primarily from in vitro reconstitution findings and observations in cultured cells(19, 22, 41, 44, 45). Further, we suggest that this may be a widespread phenomenon in cells that build elaborate actin architectures alongside microtubule reorganization.

**Figure 8:**
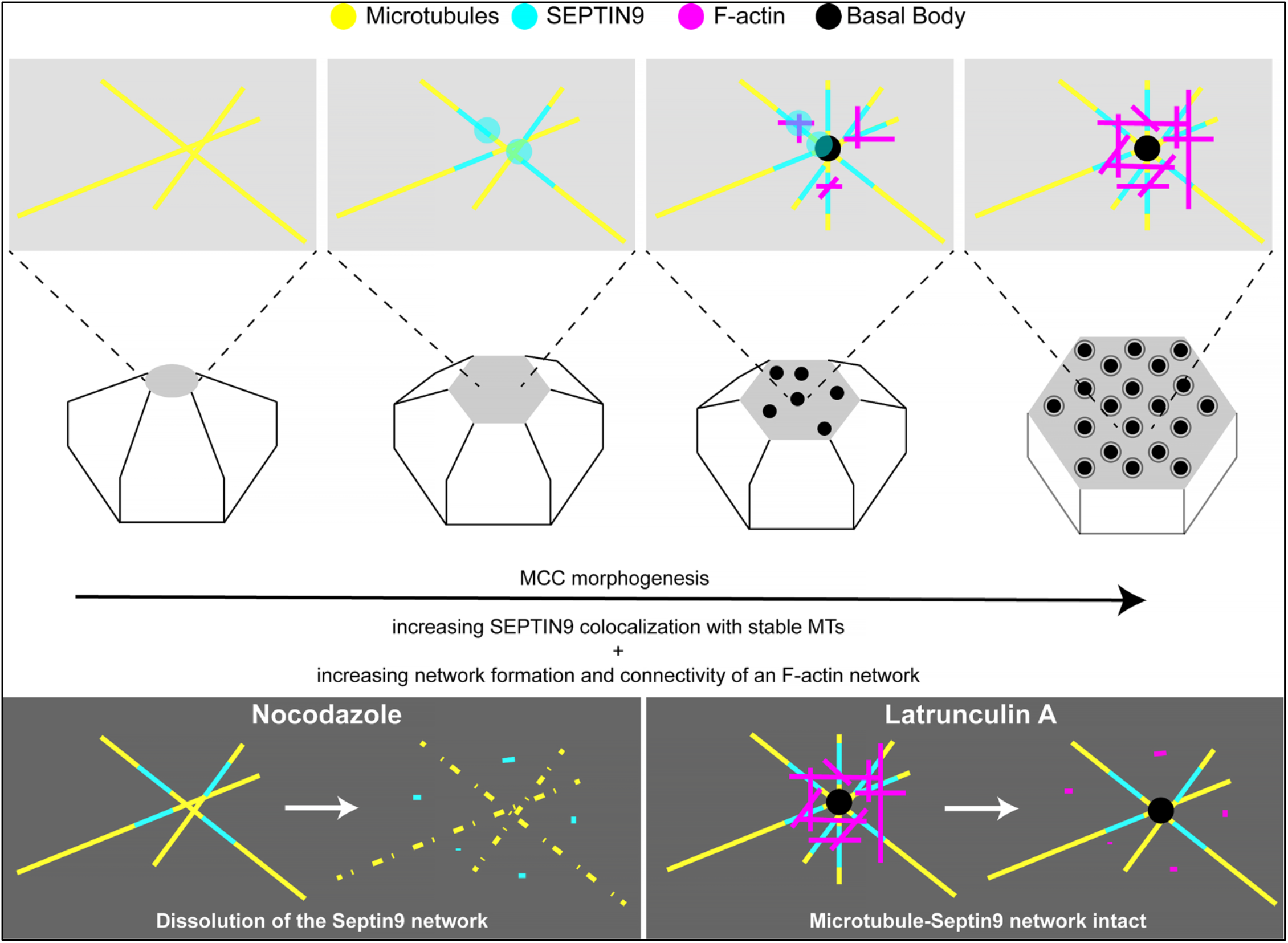
Schematic representation of the interaction between the SEPT9, microtubule, and F-actin networks. Schematic model of MCC apical expansion driven by SEPT9-mediated microtubule-to-actin remodeling. The top row shows zoomed views of the apical cytoskeleton at successive stages: microtubules (yellow) organize first, SEPT9 (cyan) assembles along them to form a mutually stabilizing scaffold, and F-actin (magenta) subsequently nucleates at microtubule-SEPT9 intersections and expands into a meshwork that scaffolds basal body docking (black). The middle row shows the corresponding apical expansion of the MCC. Bottom panels illustrate pharmacological dependencies: nocodazole disrupts microtubules and dissolves the SEPT9 network, while latrunculin A disrupts F-actin but leaves the SEPT9-microtubule scaffold intact.

Our staging analysis distinguishes MCC apical emergence from expansion: *septin9-x1* morphants are delayed in emergence but eventually reach the surface, only to fail in the post-emergence expansion phase. SEPT9 is therefore required specifically for the actin assembly that drives apical expansion, not for radial intercalation itself. This connects directly to the actin-pushing mechanism described by Sedzinski et al.(6): pushing forces require an actin meshwork, and that meshwork requires SEPT9. Whether SEP9 acts upstream of or in parallel with the RhoA-Formin 1 axis remains an open question(12). Its established interaction with the RhoGEF ARHGEF18 raises the possibility that cortical SEPT9 activates local RhoA signaling to direct formin-dependent actin nucleation(31); alternatively, SEPT9 may stabilize nascent actin filaments directly through its known crosslinking and cofilin-inhibition activities(36). Distinguishing these models will require identifying SEPT9’s binding partners at the cortical MT network.

Alternative splicing produces two SEPT9 isoforms with distinct cytoskeletal responsibilities: isoform 1 organizes both MT and actin networks via its MBD-containing N-terminal extension, while isoform 4 contributes only partially to actin through the shared acid poly-proline and GTP binding domains. The convergence of isoform 4 and ΔMBD partial-rescue phenotypes, constructs differing only in the MBD, identifies these 25 amino acids as the molecular switch separating full function from partial function. The MBD-i4 result further reveals inhibitory activity at the SEPT9 C-terminus that the N-terminal extension counteracts, providing in vivo support for the regulatory architecture proposed by Kuzmić et al., based on HNA mutations affecting MT binding(35).

A central open question is whether SEPT9 functions within the canonical SEPT2-6-7-9 hetero-octamer in MCCs or independently(26). In motile cilia, SEPT7 localizes at the base and is essential for apical F-actin enrichment, suggesting that the cortical MT scaffold described here can be assembled by an octamer(33). The main goal of future experiments will be to investigate this hypothesis in MCCs.

Our results have direct relevance for hereditary neuralgic amyotrophy (HNA), an autosomal-dominant disorder linked to SEPT9 mutations concentrated in the 25–164 region(35, 40). This same region appears to influence SEPT9 localization, shifting it from microtubules to F-actin. Consequently, HNA patients may display subtle mucociliary phenotypes. However, since HNA individuals typically retain one wild-type allele, unlike motile ciliopathies, which are generally autosomal recessive, the developmental impact of SEPT9 dysfunction may not produce overt phenotypes in these patients. Biallelic loss of SEPT9 might induce more pronounced effects; however, the embryonic lethality observed in Septin9 null mice suggests that complete loss is incompatible with life. Two approaches should provide better insight into this problem: first, assessing whether known HNA mutations induce MCC defects in our model system in an autosomal-dominant manner; second, systematically evaluating mucociliary function in HNA patient cohorts for mild abnormalities.

In conclusion, we identify SEPT9 as the molecular link that translates cortical microtubule organization into apical actin assembly during multiciliated cell morphogenesis, providing the first in vivo evidence that microtubule-to-actin coordination drives apical surface expansion.

## METHODS

### Reagents and tools table

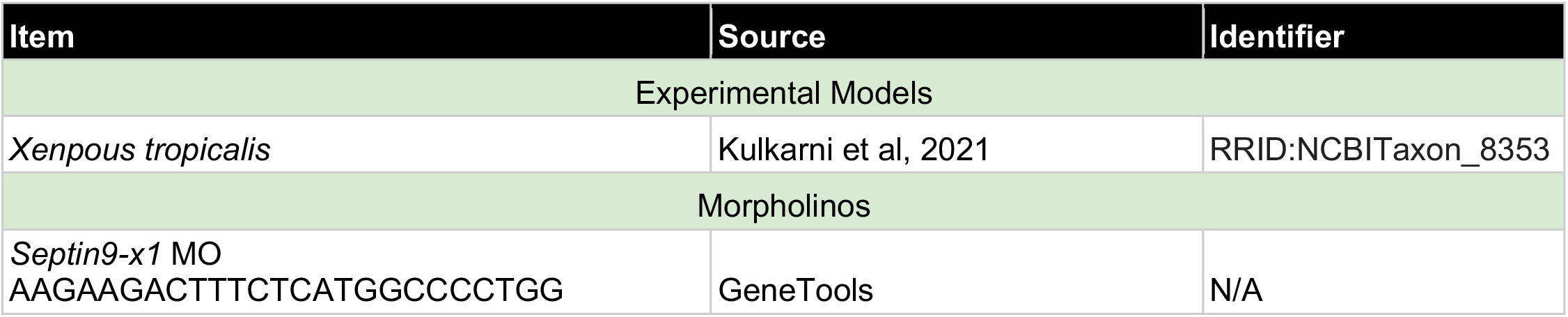
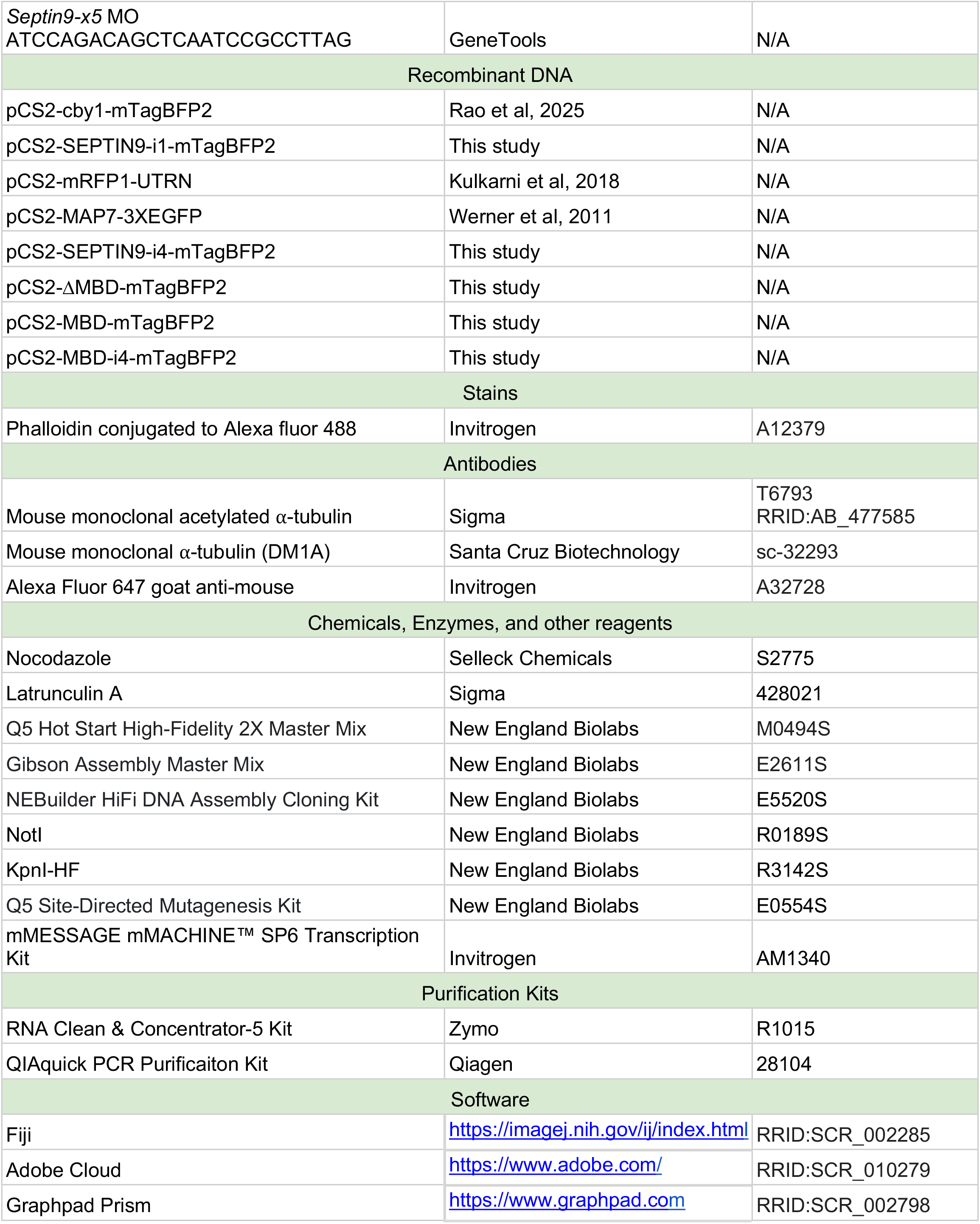

### Animal husbandry and in vitro fertilization

Frogs (*Xenopus tropicalis*) were bred and housed in a vivarium in accordance with protocols (ACUC# 4295) approved by the University of Virginia Institutional Animal Care and Use Committee (IACUC). Embryos used in experiments were generated by in vitro fertilization as described previously(46). Briefly, the testes from male frogs were crushed in 1× MBS (pH 7.4) with 0.2% BSA and added to eggs obtained from the female frogs. After 3 min of incubation, freshly made 0.1X MBS (pH 7.8) was added, and the eggs were incubated for 10 more minutes or until contraction of the animal pole of the eggs was visible. The jelly coat was removed using 3% cysteine in 1/9× MR solution (pH 7.8–8.0) for 6 min. The embryos were either microinjected with morpholino or mRNA and raised between 21-28°C in a temperature-controlled incubator (VWR, #2005) until they reached appropriate stages for experiments. Embryos were staged as described previously(47). The fertilized embryos for microinjection were chosen randomly.

### Morpholino oligos and RNA

Morpholino oligos and plasmids used in this study are listed in the reagents and tools table. Both morpholinos (septin9 X1 and septin9 X5) were designed by GeneTools. Morpholinos were resuspended to a stock solution (20µg/µL) by adding 126.5µL of pure water to 300nmol (2.53mg) of morpholino. For a working solution, morpholinos were diluted 1:4 (5 µg/µL), and 2 nL were injected per embryo, for a total of 10 ng per embryo. mRNA used in this study was generated by linearizing the sequence-verified plasmids (Not1 or KpnI with NEB CutSmart buffer), purified using the QIAquick PCR Purification Kit for PCR Cleanup (Qiagen, #28104), in vitro transcribed using the mMESSAGE mMACHINE™ SP6 Transcription Kit (Invitrogen, #AM1340), and purified using the RNA Clean & Concentrator-5 kit (Zymo, #R1015). The amount of mRNA injected varied per plasmid, as shown in the reagents and tools table. All plasmids used to generate mRNA were sequenced and verified by either Eurofins or Plasmidsaurus whole plasmid sequencing.

### Microinjections

mRNA or morpholino was injected into early-stage embryos (stage 1, 3, or 4) using 4in thin-wall glass capillaries (World Precision Instruments, #TW100F-4) pulled using a Flaming/Brown Micropipette Puller (Sutter Instruments, #P-97) to generate needles. Loaded needles were mounted to a micromanipulator (Narishige, #MM-3), and the back of the needle was fed into a Pico-Liter Injector (Warner Instruments, #PLI-10) connected to a pneumatic system to inject into early-stage *X. tropicalis* embryos. Approximately 2nL of reagent was injected into each embryo, as measured and visualized under a stereo microscope (Nikon, SMZ745).

### Microbead Flow Assay

Injected embryos were raised to stage 28 and moved to a modeling clay dish filled with 1/9× MR + 0.05% benzocaine for anesthetization (Sigma, #E1501). Then, 1µL of polystyrene microspheres (Bangs Laboratory, #DSCR006) was pipetted to the anterior end of the immobilized embryo, and images were acquired every 200ms for 10 seconds on a stereo microscope (Nikon, #SMZ745T and #DS-Fi3 camera) to visualize beads propelled from the anterior to the posterior end of the embryo. Raw data of the image sequences were extracted in Fiji and analyzed in Microsoft Excel to determine the average velocity of the leading bead front.

### Immunofluorescence staining

*X. tropicalis* embryos were fixed at the appropriate stage in 4% paraformaldehyde for 20 minutes and continuously rotated on a Belly Dancer (Stovall Life Science, #USBDbo). Then, embryos were washed 3 times in PBST (PBS + 0.2% Triton X-100) for 10 minutes each, followed by blocking (3% BSA in PBST) for 1 hour. Embryos were then incubated in primary antibody (diluted to the appropriate concentration in blocking solution) for 1 hour. Then, embryos were washed 3 times in PBST for 10 minutes, followed by incubation with secondary antibody and phalloidin (both at 1:500) for 1 hour. Lastly, embryos were washed twice with 1× PBS for 5 minutes each, transferred to a glass slide (Fisherbrand, #22339408), mounted in a single drop of mounting medium (Ibidi, #50001), and covered with a glass coverslip (epredia, #22×22-1.5-001G) for imaging. To control the pressure applied by the coverslip and avoid crushing embryos, silicone grease (Danco, #88693) was placed at all four corners of the coverslip prior to mounting. All immunostaining steps were performed at room temperature.

### Imaging

Confocal imaging was performed using the Leica DMi8 SP8 confocal microscope with a 40× oil immersion objective (1.3 NA) and 63× oil objective (1.40 NA). Images were captured between 1-12× zoom at either 512×512 or 1024×1024 resolution. Most images in this manuscript are max intensity projections of z-stacks with 0.33µM steps. Acquired images were adjusted (brightness and contrast) and analyzed in Fiji and then assembled in Adobe Illustrator software.

### Drug treatments

For microtubule disruption, stage 23 embryos (injected with SEPTIN9-i1-BFP, RFP-UTRN, and MAP7-GFP) were treated as described previously (Werner et al. 2013) with 1µM of nocodazole (Selleck Chemicals, #S2775) until stage 28. For F-actin disruption, mid-tailbud embryos were treated as described previously (Kulkarni et al. 2018) with 2µM of Latrunculin A (Sigma, #428021) for 15 minutes. DMSO-treated embryos were used as controls for both experiments. Embryos were then live mounted in 1/9x MR + 0.05% benzocaine and imaged on the Leica SP8 DMi8 confocal microscope.

### Image analysis, quantification, and statistics

All the experiments were repeated three times. All the measurements and analyses were performed on at least three embryos. Sample size, indicated by “n” values, is included in the figure legends. For statistical analysis, Prism ver. 11 was used. The type of analysis for each experiment is included in the figure legends. Fiji (ImageJ2) version 2.16.0/1.54p was used for measuring flow velocity, MCC area, basal body density, basal body uniformity, cilia abundance, colocalization, MT network connectivity, MT network complexity, F-actin network complexity, SEPT9 network complexity, MT network junctions, SEPT9 network junctions, and F-actin network junctions. Furthermore, this version of Fiji was used to deconvolve the images in figure 3 and supplemental figure 2 and to skeletonize SEPT9, MT, and F-actin networks in figure 3, figure 4, figure 5, figure 6 and supplemental figure 4. For Fiji code (groovy), ChatGPT was used to edit/streamline code. All ChatGPT-assisted code was verified before use for analysis. No artificial intelligence was used for any other part of this manuscript, including image processing and figure preparation.

## ACKNOWLEDGMENTS

We thank Dr. Karen Hirschi for providing access to the confocal microscope. We are grateful for the NIH grants: NIGMS R35GM146856.

## AUTHOR CONTRIBUTIONS

AA: Investigation, Visualization, data analysis, and manuscript writing.

VH: Investigation, Visualization

ES: Intellectual contributions and manuscript feedback.

SSK: Conceptualization, Methodology development, Supervision and mentoring, and Manuscript writing and revisions.

