## Supplemental File for "Microtubule-templated actin assembly by septin9 drives apical expansion of epithelial cells"

Supplemental Figures

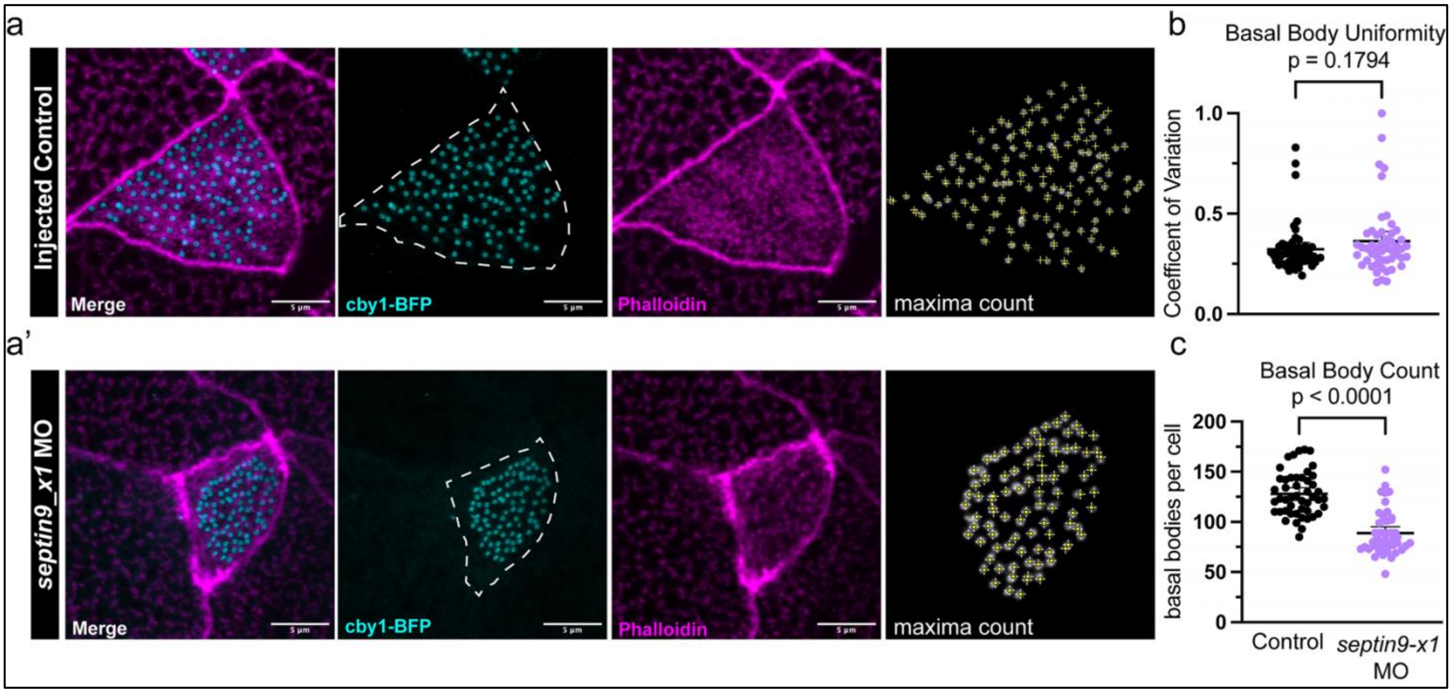


**Figure S1: Septin9 is essential for basal body number but not for uniformity in multiciliated cells.**

**(a)** Confocal max intensity projection of a cby1-BFP (basal bodies, cyan) injected stage 28 embryo stained with phalloidin (F-actin, magenta) and used to count basal bodies (maxima count) using Fiji software.

**(a')** Confocal max intensity projection of a cby1-BFP (basal bodies, cyan) injected stage 28 *septin9-x1* morphant stained with phalloidin (F-actin, magenta) and used to count basal bodies (maxima count) using Fiji software.

**(b)** Quantification of basal body uniformity determined by the coefficient of variation of nearest neighbor with Fiji software.

**(c)** Quantification of basal body number using maxima count from panel a and a'. For each analysis: n=12 embryos and 47-54 cells per condition. Scale bar is 5µm for each image.

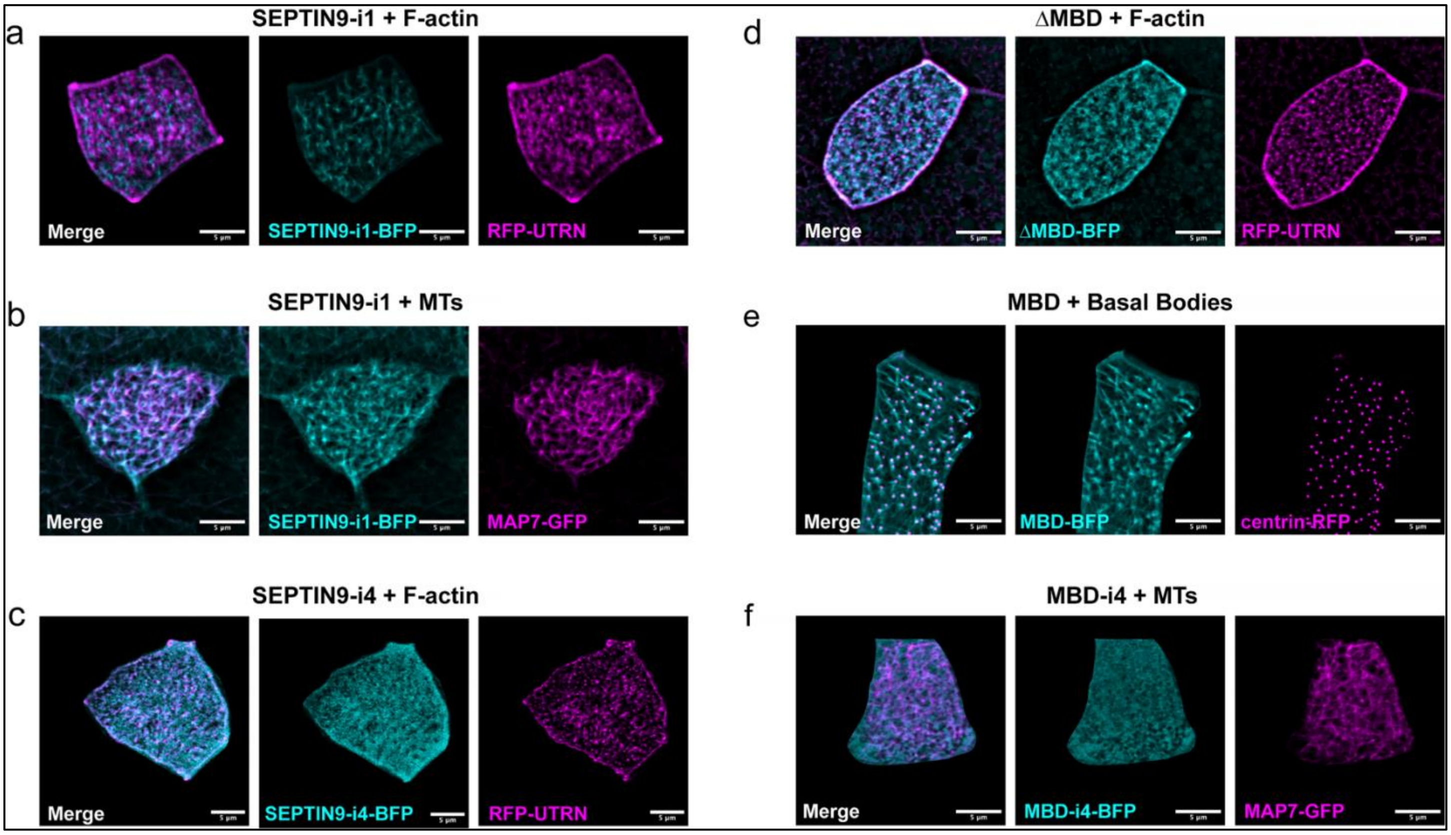


**Figure S2: Septin9 localizes to cortical microtubules, not to the F-actin network via its microtubule binding domain (MBD, 1-25 aa).**

**(a-f)** Uncropped confocal max intensity projections from Figure 3. Each image is deconvolved using Fiji software (2-5 iterations). Scale bar is 5µm for each image.

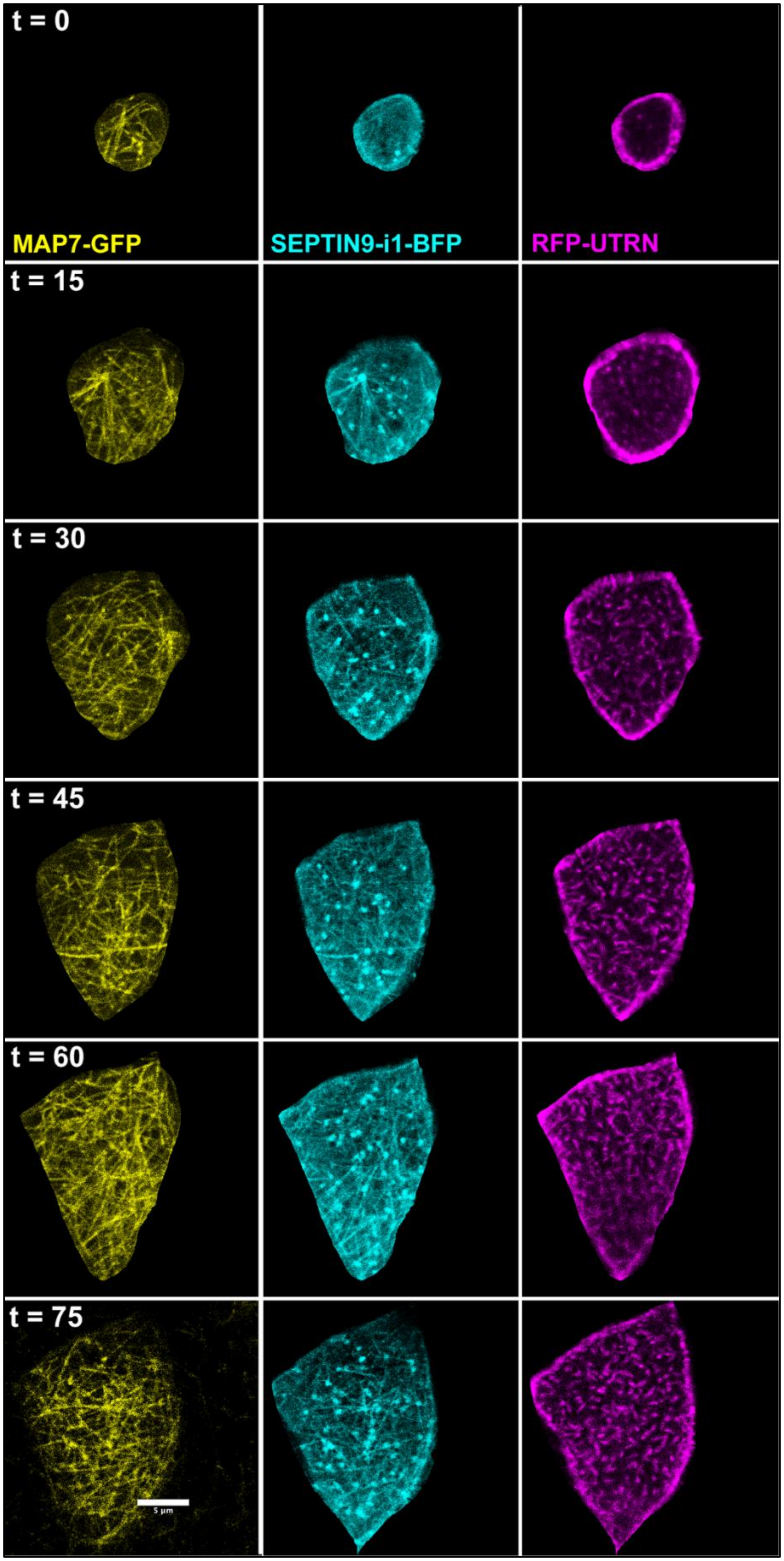
t = 0
MAP7-GFP
SEPTIN9-i1-BFP
RFP-UTRN
t = 15
t = 30
t = 45
t = 60
t = 75
5 µm

**Figure S3: Cortical microtubules template Septin9 organization followed by filamentous actin assembly.**

Each channel, MAP7-GFP (yellow), SEPT9IN9-i1-BFP (cyan), and RFP-UTRN (magenta), is shown in individual channels for the full developmental period from t=0 to t=75 minutes. Scale bar is 5μm.

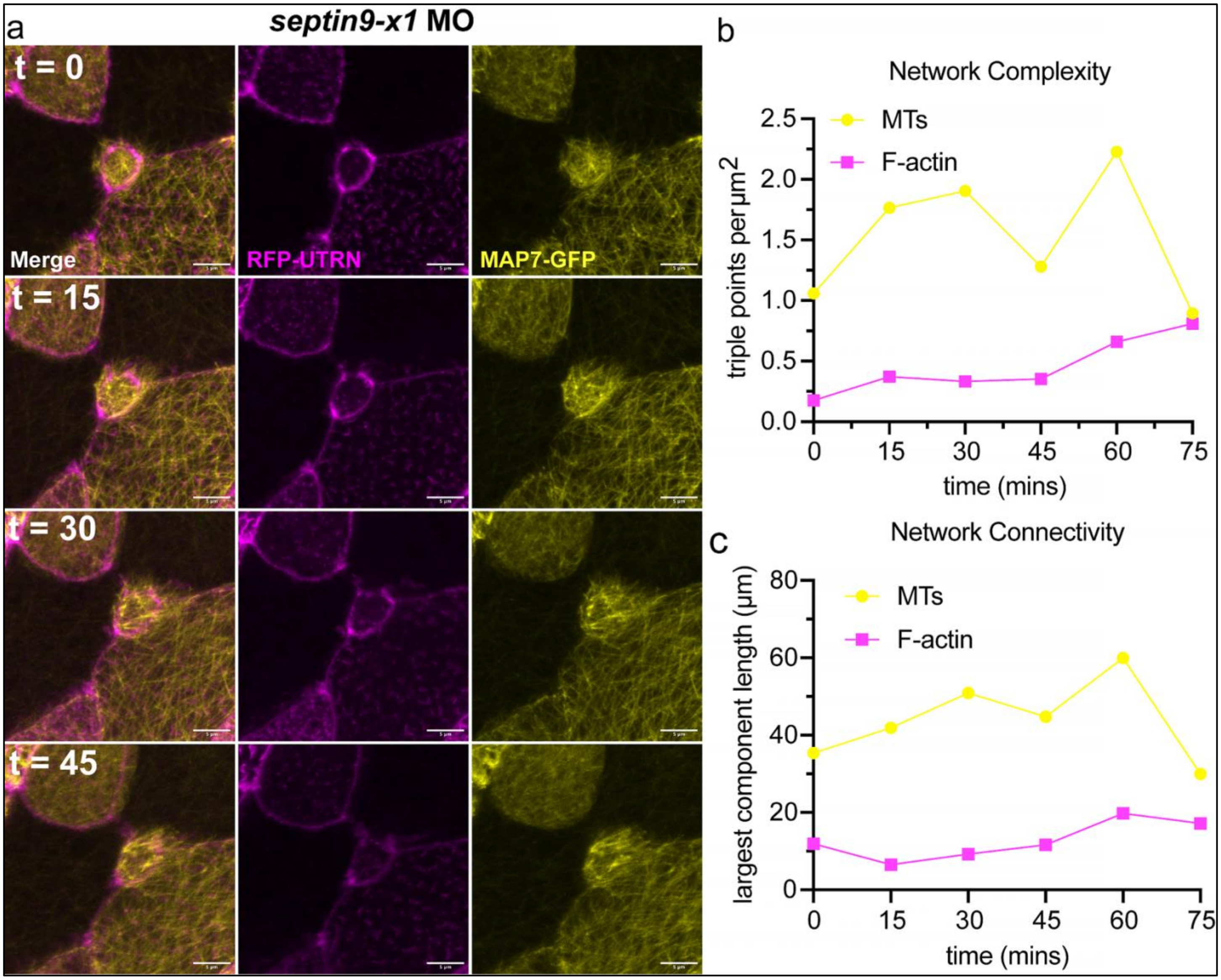


**Figure S4: Septin9 is essential for microtubule and F-actin organization in multiciliated cells.**

**(a)** Time-lapse images (confocal max intensity projections) of a newly intercalated MCC from a stage 18 *septin9-x1* morphant co-injected with SEPTIN9-i1-BFP (cyan), RFP-UTRN (F-actin, magenta), and MAP7-GFP (microtubules, yellow). The same MCC was imaged in the outer ectoderm for over 75 minutes. Only the first 45 minutes are shown as the cell neither expanded nor built an F-actin network over time.

**(b)** Quantification of the network complexity of the microtubule and F-actin networks over time from the time-lapse in panel a.

**(c)** Quantification of the network connectivity of the microtubule and F-actin networks over time from the time-lapse in panel a. Largest component length and triple points/μm$^2$ were determined from the skeletonized images of each channel using a Fiji script. Scale bar is 5μm for each image.
